# Mapping Tumor-Specific Expression QTLs in Impure Tumor Samples

**DOI:** 10.1101/136614

**Authors:** Douglas R. Wilson, Wei Sun, Joseph G. Ibrahim

## Abstract

The study of gene expression quantitative trait loci (eQTL) is an effective approach to illuminate the functional roles of genetic variants. Computational methods have been developed for eQTL mapping using gene expression data from microarray or RNA-seq technology. Application of these methods for eQTL mapping in tumor tissues is problematic because tumor tissues are composed of both tumor and infiltrating normal cells (e.g. immune cells) and eQTL effects may vary between tumor and infiltrating normal cells. To address this challenge, we have developed a new method for eQTL mapping using RNA-seq data from tumor samples. Our method separately estimates the eQTL effects in tumor and infiltrating normal cells using both total expression and allele-specific expression (ASE). We demonstrate that our method controls type I error rate and has higher power than some alternative approaches. We applied our method to study RNA-seq data from The Cancer Genome Atlas and illustrated the similarities and differences of eQTL effects in tumor and normal cells.

## 1 Introduction

Genetic variants (e.g. Single Nucleotide Polymorphisms (SNPs)) that are associated with the expression of one or more genes are referred to as gene expression quantitative trait loci (eQTLs). Genome-wide eQTL study is a powerful tool for understanding the functional roles of genetic variants. For example, eQTL analyses can help interpret the results of genome-wide association studies (GWASs) [1]. There are two types of eQTL, *cis-*eQTL and *trans-*eQTL [2, 3], which are distinguished by the pattern of expression change they induce in the genes which they affect.

To precisely define these eQTL types, we first define the term “allele”. Consider a diploid genome, which has two homologous copies of each chromosome: a maternal copy and a paternal copy. As such, each genetic locus (e.g., a SNP or a gene) has two copies within a cell, which are referred to as the two alleles of this locus. For a gene affected by a *cis-*eQTL, the expression of each allele is moderated by the genetic content of the corresponding homologous chromosome, which leads to allelic imbalance of gene expression. In contrast, for a gene affected by a *trans-*eQTL, the expression of both alleles are modified to the same extent.

The concepts of *cis-* and *trans-*eQTLs are crucial to our method development, and thus we further illustrate them by two examples. Consider a *cis-*eQTL, which is a SNP with *A* and *T* alleles. The *A* allele inhibits the binding of a transcription factor, which up-regulates the expression of a nearby gene. In contrast, the *T* allele does not affect transcription factor binding. If we refer to the two alleles of this gene by *A* or *T* allele, based on the genotype of the *cis-*eQTL on each homologous chromosome, this *cis-*eQTL leads to lower expression in the *A* allele than the *T* allele. An example of a *trans-*eQTL could be a SNP that affects the activity of a transcription factor, which in turn regulates the expression of a gene but does not preferentially bind to either allele of this gene.

*Cis-*eQTLs are often falsely conflated with local eQTL as such eQTL are often located nearby the genes they affect. *Trans-*eQTL, on the other hand, can be located anywhere in the genome in relation to the genes which they regulate [2]. It is important to reinforce that the defining characteristics of *cis-*eQTLand *trans-*eQTLare not based on their proximity to their target genes, as local eQTL can induce *cis-* or *trans-* patterns of expression change.

Traditional eQTL mapping methods implicitly assume an eQTL has the same effect on all cells within a sample. This is a reasonable assumption for samples with a relatively homogeneous cell population. However, tumor samples invariably contain both tumor cells and infiltrating normal cells (e.g. immune cells) and eQTL effects could differ between these two types of cells. To quantitatively capture this concept of inhomogeneity within a tumor cell population, we consider its tumor purity, defined as the proportion of tumor cells within the tumor sample. Previous eQTL studies in tumor samples often ignore tumor purity information and directly apply eQTL mapping methods that assume the tumor samples are composed of homogenous cells [4, 5, 6, 7, 8]. Our results show that ignoring tumor purity may lead to severely inflated type I error in the identification of tumor-specific eQTL.

In this paper, we focus on eQTL mapping using germline genetic variants. Our methods may be extended to study eQTL mapping using somatic variants, but such an extension must address the challenge of intra-tumor heterogeneity with respect to somatic mutations. To the best of our knowledge, only one previous work has considered a similar problem of cell-type-specific eQTL mapping given cell type proportion estimates [9]. Specifically, Westra et al [9] identify neutrophil-specific eQTLs using a linear model: *y* = *β*_0_ + *β*_1_*G* + *β*_2_*P* + *β*_3_*GP* where *y* is gene expression, *G* is genotype, and *P* is an estimate or proxy of neutrophil proportion in a composite tissue. Loci where eQTL effects are different between neutrophil and other cell types were identified by testing the hypothesis *β*_3_ = 0. This approach does not directly estimate or assess cell-type-specific eQTL effects. We show in our analysis that a variant of this method that explicitly models a tumor-specific eQTL effect has relatively lower power than our proposed method.

## 2 Model

Our model is an extension of the TReCASE method, which performs eQTL mapping using RNA-seq data [10]. The TReCASE method models RNA-seq data along two dimensions, Total Read Count (TReC) and Allele-Specific Expression (ASE), and simultaneously uses these two types of data for eQTL mapping [10, 3]. The TReC for a gene of interest is the total number of RNA-seq reads which map to this gene. Under the TReCASE framework, TReCs across samples are modeled by a negative binomial distribution. The ASE of a gene is quantified by the number of allele-specific reads that match the genotype of one haplotype, but not the other haplotype of this gene. Thus, an RNA-seq read is allele-specific if it overlaps with at least one SNP which is heterozygous across the two haplotypes. The number of allele-specific reads from one allele given the total number of allele-specific reads follows a beta-binomial distribution in the TReCASE framework.

The TReCASE method jointly analyzes the TReC and ASE data for *cis-*eQTL as these two types of data provide consistent information regarding the effect sizes of *cis-*eQTLs. In contrast, for *trans-*eQTL the eQTL effect is non-zero for TReC but zero for ASE, and thus only TReC data are used for mapping *trans-*eQTL. In this paper, we extend the eQTL framework of the TReCASE model for tumor eQTL studies through the incorporation of tumor purity and independent tumor- and normal-specific eQTL effects into our likelihood model. We refer to this new model as pTReCASE.

### 2.1 The Data

We assume that phased germline genotype data and RNA-seq data from tumor samples are available for *n* independent subjects. Since germline genotype data have been phased, we have genotypes for each of a subjects’ two haplotypes. We also assume that an estimate of tumor purity is available for each tumor sample. For example, one could estimate tumor purity using somatic copy number aberration data [11].

In the following discussion, we consider a specific gene of interest and a single potential eQTL of this gene. Let *G*(*i*) be the genotype of subject *i* at the potential eQTL. *G*(*i*) can take values in *{AA, AB, BB}* where A and B denote the major and minor allele of the potential eQTL. Let *ρ*_*i*_, *d*_*i*_, and ***x****_i_* = (*x*_*i1*_,…, *x*_*ip*_)*^T^* be the tumor purity estimate, read depth measurement, and a vector of *p* covariates for the *i*-th sample respectively. We set *d*_*i*_ as the 75-th percentile of the TReCs across all genes in the *i*-th sample, which is a more robust way to measure read-depth than the summation of the TReCs across all genes.

### 2.2 Total Read Count (TReC) Model

The total read count *Y*_*i*_ is defined as the number of RNA-seq reads that are mapped to a given gene. We assume that *Y*_*i*_ follows a negative binomial distribution with mean *µ*_*i*_ and over-dispersion *φ*, the likelihood for which is given by:

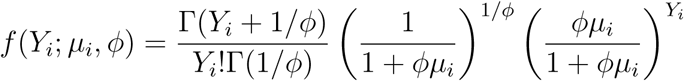

with *E*(*Y*_*i*_) = *µ*_*i*_ and 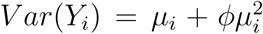. Summarizing across all *n* subjects, the log-likelihood for the TReC model is:

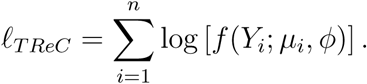

The TReC model captures the genetic effects of potential eQTL in impure tumor samples through its specification of the mean *µ*_*i*_. In order to describe this structure, we must first quantitatively define these genetic effects in both tumor and normal cells. Let *µ*_*iA*_ and *µ*_*iB*_ be the mean expression of alleles A and B for the *i*-th subject, and use superscripts ^(*T*)^ and ^(*N*)^ to denote measurements from tumor and normal cells, respectively. We define

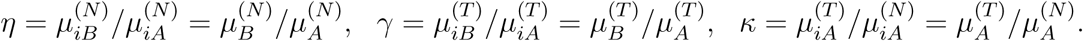

Then *η* and *γ* are eQTL effects within the normal and tumor tissues, respectively, and *κ* is a nuisance parameter that models baseline gene expression differences between tumor and normal tissues. Here we assume that *η*, *γ*, and *κ* are parameters which are shared across subjects. Let *ξ*_*i*_ = *µ*_*iB*_/*µ*_*iA*_. Assuming that the mean expression of an allele is the weighted summation of its expression in tumor and normal cells, we have:

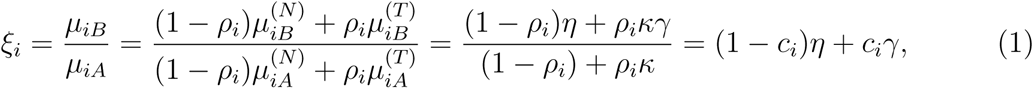

where *c*_*i*_ = (*ρ*_*i*_*κ*)/(1 *- ρ*_*i*_ + *ρ*_*i*_*κ*). The third equality is obtained by dividing both the numerator and denominator by 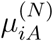. Therefore, the overall genetic effect in tumor samples is a mixture of the genetic effects within tumor cells and normal cells.

Next we consider modeling the *µ*_*i*_ across different genotypes. First, if the *i*-th subject has genotype AA at the candidate eQTL,

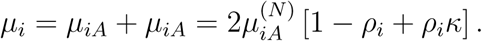

We model 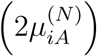 using a linear function of log read-depth and the *p* covariates: *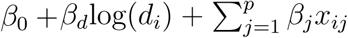*. Applying similar derivations for the subjects with genotypes AB and BB, we have:

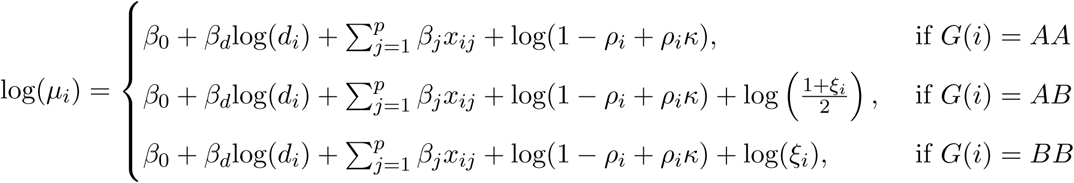

Model fitting is accomplished via block coordinate ascent using two blocks: block 1 consists of *κ, η,* and *γ* and block 2 consists of *φ*, *β*_*d*_, *β*_*j*_ for *j* = 0, 1, *…, p*. Given the initial values of the parameters in block 2, the parameters of block 1 are updated via a quasi-Newton method (LBFGS). Then, given the parameters in block 1, the parameters in block 2 are updated via negative binomial regression. We iteratively update the parameters in blocks 1 and 2 until convergence.

### 2.3 Allele Specific Expression (ASE) Model

We refer the reader to Sun and Hu [3] for details on how AS reads are counted in RNA-Seq data. In the following, we briefly describe this process for a single candidate eQTL. For each subject, we have genotypes available for arbitrarily labeled haplotypes, haplotype 1 and haplotype 2. We extract all RNA-seq reads that overlap with at least one heterozygous SNP within the body of the gene and assign each of these reads to the haplotype which matches its nucleotide sequence. As haplotypes 1 and 2 are arbitrarily labeled for each subject, we ensure comparability across subjects by relabeling these haplotypes with respect to the genotype of the candidate eQTL. For subjects who are heterozygous at the candidate eQTL, haplotype *A* contains the *A* allele of the candidate eQTL and haplotype *B* contains the *B* allele. For subjects who are homozygous at the candidate eQTL, haplotypes *A* and *B* may be defined arbitrarily without affecting the likelihood function or statistical inference.

Let *R*_*iA*_ and *R*_*iB*_ be the number of allele-specific RNA-seq reads assigned to haplotypes *A* and *B*, respectively. Let *R*_*i*_ = *R*_*iA*_ + *R*_*iB*_ be the total number of allele-specific RNA-seq reads. We model *R*_*iB*_ given *R*_*i*_ using a beta-binomial distribution with probability of success *π*_*i*_ and over-dispersion *ψ*, the likelihood of which is given by:

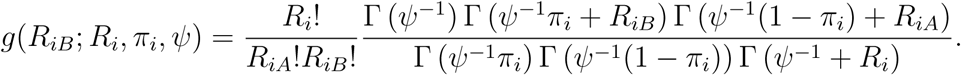

Incorporating all individuals, we may express the log-likelihood for the ASE model as:

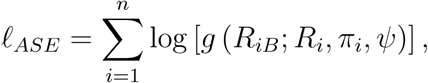

where

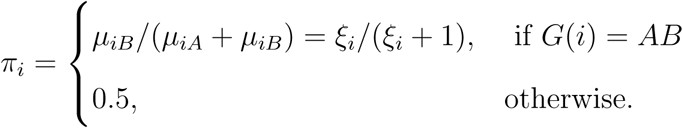

A consequence of the above modeling strategy is that subjects who are heterozygous at the potential eQTL do not contribute to the estimation of the eQTL parameters *κ, η*, or *γ*. However, such subjects are informative regarding the over-dispersion parameter *ψ*.

Model fitting is achieved via block coordinate ascent using two blocks: block 1 consists of *κ*, *η* and *γ*; block 2 consists of *ψ*. We employ a cyclical algorithm similar to the one used in the previous section, except that LBFGS is used to update the parameters of both blocks.

### 2.4 TReCASE: Unifying TReC and ASE Models

Restricting to *cis-*eQTLs, the TReC and ASE models share the *κ*, *η*, and *γ* parameters allowing for unification into a single likelihood model:

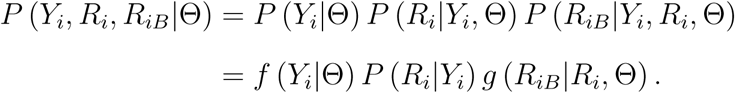

Note that the relationship above explicitly relates *Y*_*i*_ and *R*_*i*_ since *R*_*i*_ ≤ *Y*_*i*_. It is reasonable to assume that given *Y*_*i*_, the distribution of *R*_*i*_ does not depend on our covariate or eQTL effects. Rather, its distribution is a function of the number of heterozygous SNPs present within the gene-body. Therefore, we may remove *P* (*R*_*i*_|*Y*_*i*_) from the likelihood function. The log-likelihood of all *n* subjects is then given by:

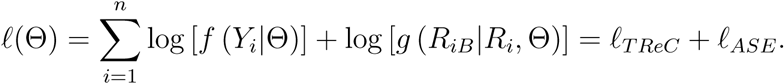

Model fitting is achieved via block coordinate ascent using three blocks: block 1 consists of *κ*, *η* and *γ*; block 2 consists of *φ*, *β*_*d*_ and *β*_*j*_ for *j* = 0, 1, *…, p*; and block 3 consists of *ψ* alone. A single update is defined by the following steps. First, given the parameters of blocks 2 and 3, the parameters of block 1 are updated using LBFGS. Then, given the parameters of blocks 1 and 3, the parameters of block 2 are updated via negative binomial regression. And finally, given the other parameters, the parameter of block 3 is updated using LBFGS. These cyclical updates are repeated until convergence.

### 2.5 Hypothesis Testing

Under the proposed models of sections 2.2 through 2.4, there are three critical questions of interest. Does an eQTL exist within normal tissue? Does an eQTL exist within tumor tissue? Should we use the TReC or TReCASE model to detect an eQTL?

The presence of eQTL in normal tissue (i.e., *η ≠* 1) or tumor tissues (i.e., *γ ≠* 1) can be assessed using likelihood ratio tests (LRT):

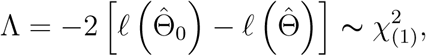

where 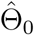 represents parameter estimates under the null and 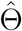 represents estimates under the alternative. To test for the presence of an eQTL in normal or tumor tissue, 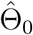 is obtained by fitting the model under a null hypothesis of *η* = 1 or *γ* = 1, respectively.

Addressing the final question requires consideration of the biological mechanisms driving *cis-* and *trans-*eQTLs. For a *cis-*eQTL, the TReC and ASE components share the same parameters for eQTL effect sizes, and thus jointly modeling TReC and ASE (i.e., TRe-CASE) increases power. For a *trans-*eQTL, expression of both alleles of the affected gene are altered to the same extent, and thus ASE is not informative in the detection of eQTL or estimating eQTL effect size. Therefore, only TReC data should be used for eQTL mapping of *trans-*eQTL. We develop a “Cis-Trans” score test to aid in model selection by addressing a null hypothesis of consistent eQTL effects across the TReC and ASE components of the model.

To structure this test, let *η*_*ASE*_ and *γ*_*ASE*_ be the eQTL effects for a gene and a candidate eQTL within the ASE component of the model. We still use *η* and *γ* to model eQTL effects in TReC data. Define *η*_*ASE*_ = *η* + *α*_*η*_ and *γ*_*ASE*_ = *γ* + *α*_*γ*_ where *a*_*η*_ and *a*_*γ*_ reflect the discrepancy between ASE and TReC eQTL effects for both normal and tumor tissues, respectively. The null hypothesis of equivalent eQTL effects in TReC and ASE components of the model is defined using the notation above by *a*_*η*_ = *a*_*γ*_ = 0. See the supplementary information for a detailed description and derivation of this “Cis-Trans” score test. The test statistic and its asymptotic distribution are provided below:

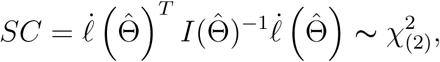

where 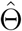 are the MLEs of our parameters under the null hypothesis; 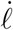 is the gradient of the TReCASE likelihood with respect to the parameters; and 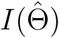 is the Fisher’s Information Matrix.

## 3 Results

### 3.1 Simulation Study

We conducted a simulation study to compare the statistical power and type I error rate of pTReCASE and several other methods. Simulations were conducted across a range of eQTL effect sizes in normal and tumor cells. To assess Type I error, we set *γ* at 1 and allowed *η* to vary. To assess power to detect tumor-specific eQTL, we set *η* at 1 and allowed *γ* to vary. For each pair of *η* and *γ*, we simulated 400 replicates of gene expression and genotype data for 500 subjects. Genotypes were simulated assuming a minor allele frequency of 0.2. Read counts were simulated according to the pTReCASE model using the following algorithm:

(1) Randomly generate tumor purities from a uniform distribution on (0.5,1) for each of the 500 subjects.

(2) Simulate TReC via a negative binomial model with:

(A) Mean of 100 reads for subjects with genotype AA and tumor purity of 0%.

(B) *κ* = 1.5 and *φ* = 0.2.

(3) Assume that 5% of the simulated TReC reads are AS, rounded to the nearest integer. Partition AS reads to haplotypes according to the established beta-binomial model using an overdispersion of *ψ* = 0.2.

Each considered eQTL model is then fit to the simulated data. For any given modeling procedure, the type I error is estimated by the proportion of simulations which incorrectly identify a tumor eQTL when none is present. Power is estimated by the proportion of simulations which correctly identify a tumor eQTL when one is present.

The competing eQTL models that we considered include the TReC/TReCASE model without correction for tumor purity, and the TReC model with tumor purity (pTReC). In addition, we also considered a naive approach of linear regression ignoring tumor purity, labeled LR, and a modification of the approach adopted by Westra et al [9] denoted by pLR. To fit a linear model, we first applied a normal quantile transformation to (read-depth corrected) TReC values of each gene across *n* samples, and then used the transformed TReC as a response variable for linear regression. Specifically, we first replaced TReC values by their ranks across *n* samples, and then replaced the ranks by their corresponding normal quantiles. For example, rank *r* was replaced by the *r/*(*n* + 1)-th normal quantile. Letting 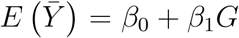 be the transformed TReC data, the linear model is given by 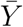, where *G* is the genotype of the candidate eQTL.

To test genotype and tumor purity interaction using pLR, we fit a linear model 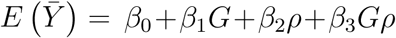 where *ρ* is an estimate of tumor purity. The interaction test employed by Westra et al [9] (i.e. *β*_3_ = 0) does not assess the strength of a tumor eQTL. Rather, it tests whether eQTL effects differ between tumor and normal tissues. Under pLR, we assessed tumor eQTL effects by testing *β*_1_ + *β*_3_ = 0 since *β*_1_ + *β*_3_ is the genetic effect of the candidate eQTL when tumor purity is 1.

All three methods that control for tumor purity (pTReCASE, pTReC, pLR) control Type I error at the desired level. As eQTL strength in the normal tissue increases, the methods that do not account for tumor purity see a rapid increase in Type I error (Figure 1A). In terms of power (Figure 1B), the methods that do not account for tumor purity exhibit the largest statistical power due to their anti-conservative control of Type I error. Among those methods that control Type I error (i.e. pLR, pTReC, pTReCASE), pTReCASE has the highest power. This is a result of its joint analysis of TReC and ASE.

**Figure 1.**
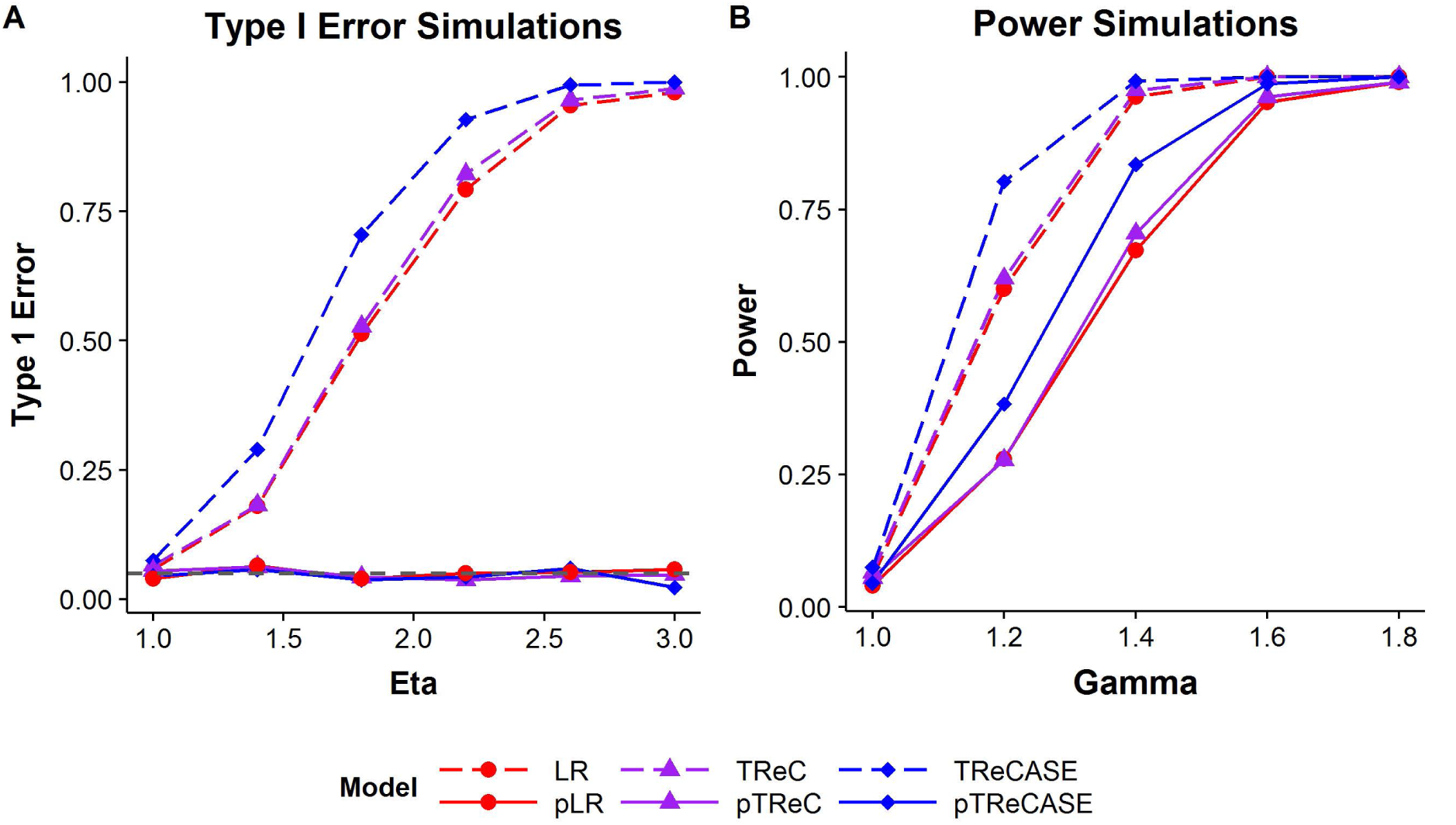
Examining Type I error [A] and Power [B] from simulation studies.

### 3.2 The Cancer Genome Atlas (TCGA) Data

We applied the pTReCASE model to analyze gene expression and germline SNP genotype data from 550 breast cancer patients of The Cancer Genome Atlas (TCGA) project. We started with 728 patients with RNA-seq data from tumor samples. In order to assess allele-specific gene expression, we downloaded raw RNA-seq data in bam file format. For genotype data, we downloaded the Affymetrix CEL files. We restricted our analysis of these 728 patients to 550 subjects who had available genotype data, passed quality controls for both genotype and RNA-seq data, and were Caucasian females (See Supplementary Materials Section B for details). Males were excluded as breast cancer in men are rare and may have a different disease etiology from those found in women. The restriction to Caucasian samples is not necessary, but it helps to eliminate possible confounders [12].

For the remaining 550 subjects, genotype imputation and haplotype phasing was performed by MACH [13] using reference haplotypes from the 1000 Genomes Project. Starting with *∼*800,000 SNPs genotyped by Affymetrix 6.0 array, we imputed the gneotypes for *∼*36 million SNPs. For each sample, we used all the SNPs with heterozygous genotypes to estimate allele-specific expression (See Supplementary Materials Section B for details). For the purposes of eQTL mapping, we restricted our analysis to those SNPs with MAF *≥* 0.02 (6,825,065 SNPs after imputation) because there is limited power to detect eQTL at lower values of MAF. Tumor purities were estimated using ABSOLUTE [11], which led to the exclusion of three additional subjects lacking valid purity estimates. Estimated haplotypes and tumor purities were treated as truth in the subsequent pTReCASE and linear regression models.

Linear models for eQTL analysis and the revised Westra approach (i.e. pLR) were fit using matrixEQTL [14] and customized R code on normal quantile transformed RNA-Seq count data. TReC, TReCASE, pTReC and pTReCASE models were fit using our own package. The median analysis time for a single gene-SNP pair using pTReCASE was 2.71 seconds (IQR = 2.93 seconds) with a maximum time to solution of 7 minutes. The covariates used for eQTL mapping include read-depth of RNA-seq experiments (Supplementary Figure 7), RNA sample plates, age, and the top two principal components derived from the genotype data of the 550 Caucasian samples. Since our method is designed to identify *cis-*eQTLs and most *cis-*eQTLs are local to the genes which they affect, we restricted our analysis to SNPs located within 100Kb of the gene of interest.

Figures 2A-B illustrate a tumor-specific eQTL identified by the pTReCASE model. The estimates of effect sizes (ratio of gene expression of the *B* allele versus the *A* allele) for normal and tumor-specific eQTLs are 0.96 (*η*) and 3.51 (*γ*), respectively. The fold change of gene expression in tumor versus normal cells (for genotype AA) is 0.19 (*κ*) (Figure 2D). In other words, gene expression in tumor cells is lower than that in normal cells, but the eQTL effect is only present in tumor cells. These numerical estimates were well demonstrated by Figures 2A-B. As tumor purity increases, gene expression measured by TReC decreases (Figure 2A), and the strength of the eQTL increases. Both TReC and ASE show consistent signals that the *B* allele has higher expression, with a “Cis-Trans” test p-value of 0.95.

**Figure 2.**
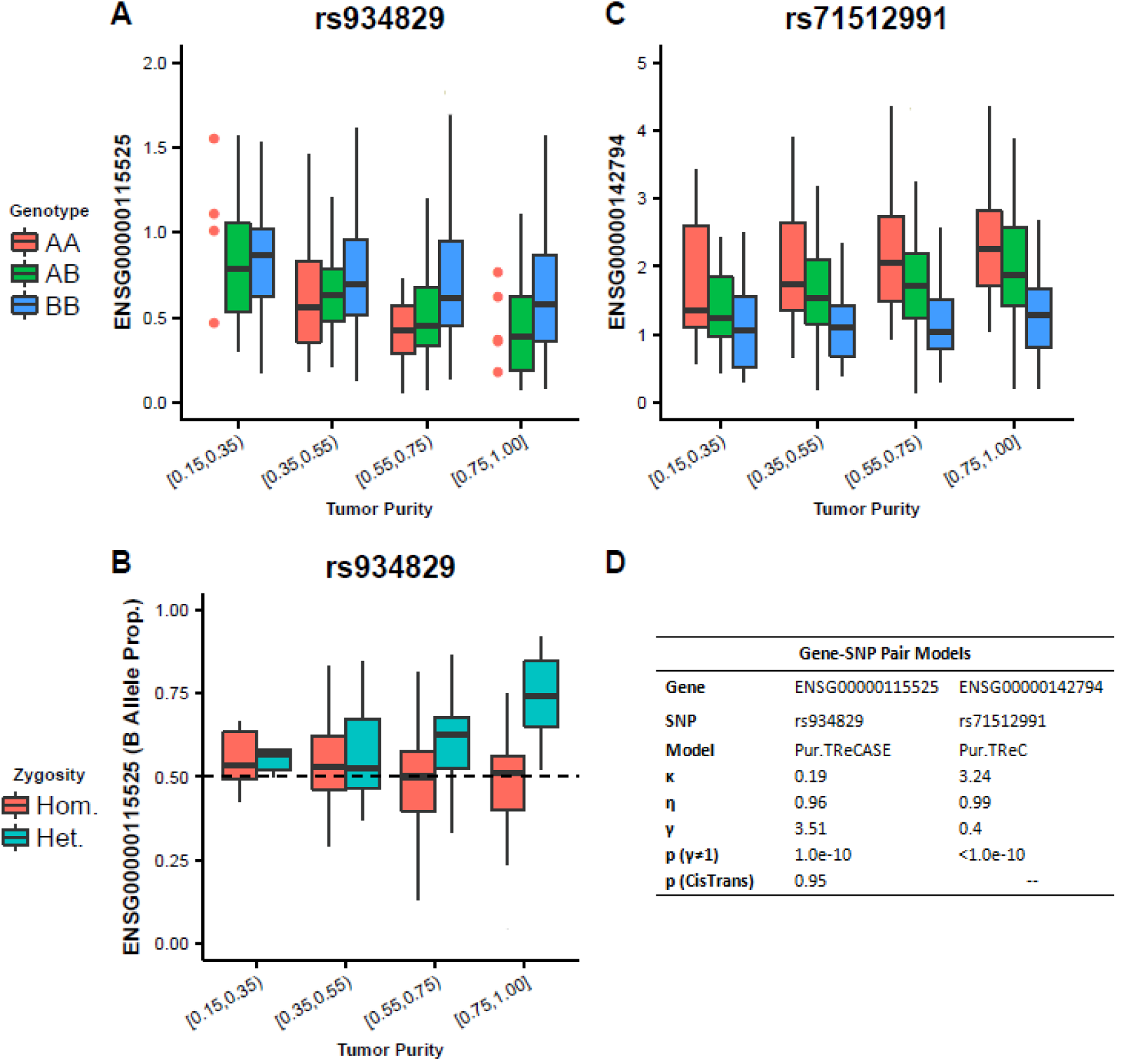
(A) Covariate-corrected total expression estimated via pTReCASE plotted against genotype and tumor purity. Outliers were suppressed for clarity. Dot plot instead of boxplot was used when sample size of a category is too small. (B) Examination of the allele specific expression corresponding to case shown (A). (C) Covariate-corrected total expression estimated via pTReC plotted against genotype and tumor purity. (D) Table providing Gene, SNP, and estimated parameters for the displayed assessments.

To highlight the functioning of the pTReC model, another example of tumor-specific eQTL was identified and shown in Figure 2C. In this example, gene expression from the ASE model is not fit due to a significant “Cis-Trans” test using the full model. In this example, gene expression is higher in tumor compared to normal cells and the *B* allele has lower expression than the *A* allele in tumor cells, but not in normal cells. Note that we can still see some signals of an eQTL in the category with the lowest tumor purity. This results from TCGA samples being selected to have relatively higher tumor purity, thereby creating a categorization schema wherein even the lowest tumor purity category has a non-negligible amount of tumor cells.

We use another example to demonstrate the utility of the Cis-Trans score test (Figure 3). Considering only the TReC data, the *B* allele has slightly higher expression than the *A* allele when tumor purity is high (Figure 3A). In contrast, considering only the ASE data, the *B* allele has much lower expression than *A* across all tumor purity levels. This inconsistency between TReC and ASE data led to a highly significant Cis-Trans p-value (Figure 3C). In such cases, only the TReC data is trusted and used to estimate eQTL effects. ASE tends to be noisier in real data as mapping biases, incorrect genotype data, and/or other biological and technical factors can lead to the observed ASE imbalance as opposed to eQTL effects. Failure to consider the Cis-Trans test could lead to the acceptance of spurious eQTL results.

**Figure 3.**
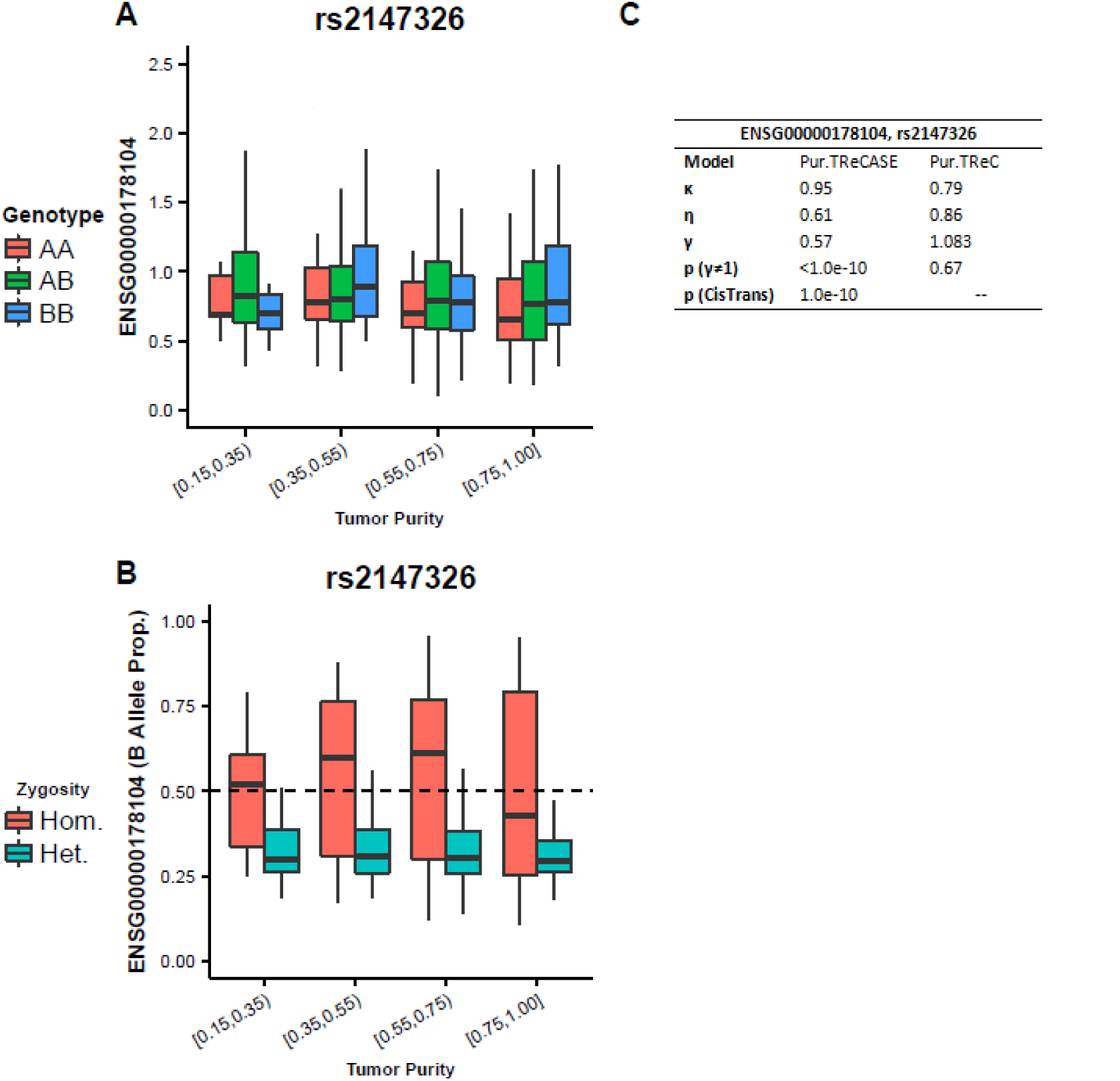
Demonstrating the utility of the Cis-Trans score test. (A) Covariate-corrected total expression plotted as a function of genotype and tumor purity. (B) Allele Specific Expression with respect to genotype and tumor purity. (C) Table containing relevant modeling information for A and B.

Next, we systematically compare the results for all eQTLs using the pTReCASE, TRe-CASE, and pLR approaches at various p-value cutoffs. One way to compare the results is to check the overlap of each significant eQTL association, i.e. each gene-SNP pair (Supplementary Table 2). However, due to LD, the expression of one gene may be associated with multiple SNPs that are in close proximity to one another and often represent redundant eQTL signals. Therefore, we focus on the eQTL results summarized at the gene level. In other words, for a given p-value cutoff, we count the number of genes with at least one eQTL with a p-value falling below the cutoff (Table 1).

**Table 1:** Summarizing the results of pTReC(ASE), TReC(ASE), the Westra-inspired models for TCGA data. Here the notation pTReC(ASE) indicates that we used the pTReCASE or pTReC model depending on the results of the Cis-Trans test. “Overlap” represents the genes with at least one significant eQTL identified by both pTReC(ASE) and an alternative method. “Overlap/alternative” is the number of overlaps divided by the number of findings using the alternative method. “Overlap/pTReC(ASE)” is the number of overlaps divided by the number of findings using pTReC(ASE). If we consider the results of pTReC(ASE) as true findings, then “overlap/alternative” is the true discovery rate and “overlap/pTReC(ASE)” is the sensitivity

| Genes |  |  |  |  |
| --- | --- | --- | --- | --- |
| P-value Cutoff | Category | Pur.TReC(ASE) | TReC(ASE) | pLR |
| $5 \times 10^{-4}$ | # of Genes | 4816 | 7793 | 2055 |
|  | overlap/alternative | – | 42.8 | 70.4 |
|  | overlap/pTReC(ASE) | – | 69.3 | 30.0 |
| $5 \times 10^{-6}$ | # of Genes | 1271 | 2982 | 268 |
|  | overlap/alternative | – | 26.8 | 84.7 |
|  | overlap/pTReC(ASE) | – | 62.8 | 17.9 |
| $5 \times 10^{-8}$ | # of Genes | 518 | 1612 | 110 |
|  | overlap/alternative | – | 21.3 | 94.5 |
|  | overlap/pTReC(ASE) | – | 66.4 | 20.1 |

Compared to pTReCASE, the TReCASE model identifies eQTLs in a larger number of genes. For those genes where TReCASE identifies a significant eQTL and pTReCASE does not, the significant findings of the TReCASE model are most likely driven by an eQTL in normal tissue. TReCASE recaptures around two-thirds of eQTL findings identified by pTReCASE. The one-third missed by TReCASE are more likely to have weaker effect size and/or are only present in tumor cells. Across p-value thresholds, the pLR model identifies relatively fewer significant gene-SNP pair relationships. Of those relationships identified by the pLR model, 70-to 90-percent are also identified by pTReCASE. The pLR model also misses at least 70% of significant results identified by pTReCASE. Possible reasons for the poor performance of the pLR model could arise from the fact that ASE is not incorporated and/or the relationship between transformed gene expression and tumor purity is not examined on the linear scale.

## 4 Discussion

Due to contamination of tumor samples with infiltrating normal cells, the identification of eQTL within tumor tissues poses several challenges. First and foremost, one needs to separately estimate the eQTL signals in tumor and normal cells. Second, while total gene expression has been widely used for transcriptome studies, it is important to leverage the additional information provided by allele-specific expression which can be effectively derived using RNA-seq data. We have developed a statistical model and software package, pTReCASE, to address these issues. The desirable performance of pTReCASE has been validated using simulations and real data analysis. In constrast, a naive approach for eQTL mapping that ignores tumor purity may lead to a large fraction of false positives.

Within the current established framework, there are three notable avenues for further development and research. The first is to improve the computational efficiency of our software package. Using the current implementation, it takes about 10,000 CPU hours for genome-wide local eQTL mapping. This can be easily done using a moderately sized computing cluster, but is not computationally feasible for a single computer. High computational costs also prevent us from using permutations to assess the significance of eQTL results. Thus, we recommend use of Bonferroni correction, Benjamini-Hochberg FDR control [15], or calculation of the number of independent tests by examining the correlation structure of the genotype data [16].

We have assumed that the haplotypes connecting candidate eQTLs and the SNPs within the gene body are known. In practice, such haplotypes are imputed/phased using statistical methods. Phasing is usually accurate within short genetic distances around the gene of interest. However, if we would like to consider potential eQTLs further from the gene, there is a possibility of phasing error. The second avenue for improving the posited model is to allow for uncertainty in the haplotype phasing by following the approach of Hu et al [17].

Lastly, both the TReC and ASE components of the pTReCASE model have assumed no copy number change across subjects. Suppose that an eQTL for a given gene modifies expression through copy number changes. More specifically, suppose the *B* allele at the candidate eQTL is associated with a larger copy number in tumor tissues. This would lead to increased expression from the *B* allele and would be interpreted by the pTReCASE model as higher expression of a single *B* allele. In such a scenario, the pTReCASE model is applicable without modification. However, if both eQTL and copy number affect gene expression independently, the pTReCASE model should be adjusted for copy number differences. Given estimates of allele-specific copy number [18], our model can be modified to address copy number variation across subjects. However, estimation of allele-specific copy number in tumor samples is a very challenging task due to the confounding effects of tumor purity, ploidy, and the possibility of subclonal copy number changes [19]. It is desirable to systematically study the effects of both germline SNPs, somatic copy number changes, and even somatic point mutations (single nucleotide variants or indels) while also accounting for intra-tumor heterogeneity, but such explorations are beyond the scope of this paper and warrant a series of future studies.

## SUPPLEMENTARY MATERIAL

**Supplement to “Mapping Tumor-Specific eQTLs in Impure Tumor Samples”:** Supplementary document containing RNA-seq and genotype array processing information, mathematical details for the optimization of the pTReC and pTReCASE models, and the derivation of the Cis-Trans score test. (PDF)

**pTReCASE:** R-package pTReCASE containing code to perform the pTReCASE analysis presented in the simulation studies and TCGA Data examination. (GNU zipped tar file)

**TCGA Data Set:** Data set used in the illustration of pTReCASE. (.txt file)

